# 5-HT1B receptors mediate dopaminergic inhibition of vesicular fusion and GABA release from striatonigral synapses

**DOI:** 10.1101/2024.03.14.584991

**Authors:** Maya Molinari, Ori J. Lieberman, David Sulzer, Emanuela Santini, Anders Borgkvist

## Abstract

The substantia nigra pars reticulata (SNr), a crucial basal ganglia output nucleus, contains a dense expression of dopamine D1 receptors (D1Rs), along with dendrites belonging to dopaminergic neurons of substantia nigra pars compacta. These D1Rs are primarily located on the terminals of striatonigral medium spiny neurons, suggesting their involvement in the regulation of neurotransmitter release from the direct pathway in response to somatodendritic dopamine release. To explore the hypothesis that D1Rs modulate GABA release from striatonigral synapses, we conducted optical recordings of striatonigral activity and postsynaptic patch-clamp recordings from SNr neurons in the presence of dopamine and D1R agonists. We found that dopamine inhibits optogenetically triggered striatonigral GABA release by modulating vesicle fusion and Ca^2+^ influx in striatonigral boutons. Notably, the effect of DA was independent of D1R activity but required activation of 5-HT1B receptors. Our results suggest a serotonergic mechanism involved in the therapeutic actions of dopaminergic medications for Parkinson’s disease and psychostimulant-related disorders.

## Introduction

Dopamine (DA) exerts a powerful influence on basal ganglia circuit activity, enabling essential motor-related functions including action selection, learning, and habit formation. It is widely recognized that the effects of endogenous DA release on motor behavior, as well as the therapeutic efficacy of medications targeting DA receptors such as levodopa and psychostimulants, originate from the activation of DA receptors within the striatum ^1^, the primary input nucleus of the basal ganglia. DA receptors exist in additional regions of the basal ganglia ^2^, although the understanding of their involvement in behavior remains limited ^3, 4^.

Particularly high levels of DA D1 receptors (D1R), the most abundant DA receptor type in the mammalian brain, are found within the substantia nigra pars reticulata (SNr) ^5, 6^, one of the output nuclei of the basal ganglia. Ultrastructural studies indicate that these D1Rs are located presynaptically on the axons of the GABAergic striatal medium spiny neurons (MSNs) enriched in D1R-expression ^7–9^. While the actions of DA on D1-MSN cell bodies and dendrites have been extensively studied, the role of DA in the regulation of neurotransmission from D1-MSN axons remains less clear. In the MSN cell body, stimulation of striatal D1Rs, which are G_olf_-coupled receptors ^10^, activates adenylyl cyclase, thereby stimulating the cAMP pathway and protein kinase A (PKA)-mediated phosphorylation of cAMP-regulated phosphoprotein of 32kDA (DARPP-32), a critical integrator of DA signaling ^11^. This cascade of molecular events results in DA-dependent regulation of gene transcription, neuronal excitability and synaptic integration, which are processes critically involved in synaptic plasticity ^11^. In contrast, the SNr has relatively little G_olf_ ^10^ or DA-sensitive adenylyl cyclase ^12^, but maintains high levels of D1R and DARPP-32 ^12, 13^. These reports suggest that the molecular machinery of somatic D1R on striatal MSN cell bodies might differ from that of presynaptic D1Rs on striatonigral axons.

DA is released in the SNr from the dendrites of substantia nigra pars compacta neurons ^14^. Thus, local DA could act on D1Rs located on the striatonigral axons and modulate GABA release, thus influencing the inhibition of the postsynaptic cells in the SNr. Most SNr cells are tonically active GABAergic projection neurons ^15^, that constitute the fundamental inhibitory unit in the disinhibition-model for movement control by the basal ganglia ^16^. Consequently, presynaptic D1Rs are in a strategic position to control motor activity by regulating the strength and timing of inhibition in the SNr, which will influence volitional movement control. Indeed, local stimulation of D1Rs in the SNr of rodents produces robust effects on behaviors such as rotation and spontaneous locomotion ^17, 18^. However, lower expression of canonical downstream components of the D1R signaling pathway suggests that DA may act via alternative mechanisms on D1-MSN axon terminals in SN as well.

Despite research interest over many years ^18–24^, the function of the D1R within in the SNr remains unclear. In part, this is because of technical challenges in isolating its effect on the local synaptic circuitry of the SNr. Many of these studies investigated the effects of DA and D1Rs with indirect methods, such as postsynaptic recordings of GABAerigc SNr cells. However, in addition to striatonigral inputs, SNr cells receive GABAergic input from the globus pallidus externa via pallidonigral projections ^25, 26^ and collateral inputs from GABAergic SNr neurons, which are tonically active ^15^ even in brain slice preparations ^27^.

To investigate the effect of DA and presynaptic D1Rs on GABA release in the SNr, we used two-photon microscopy to image synaptic vesicle fusion at the striatonigral synapses in brain slices and measured striatonigral GABA_A_ currents in SNr neurons using optogenetics. Our findings indicate that D1Rs do not affect GABA release from striatonigral synapses, but that DA potently inhibits striatonigral GABA release through serotonergic 5-HT1B receptors, consistent with a recent report in striatum ^28^.

## Results

### Labeling of D1-MSN boutons with FM1-43

D1-MSNs give rise to a prominent bundle of axons that exits from the striatum and terminates within the midbrain. A significant proportion of this projection can be preserved in sagittal mouse brain sections, and electrical stimulation of this fiber tract evokes distal GABA release from D1-MSN synapses in the SNr. We took advantage of this preparation to introduce optical recordings of synaptic vesicle fusion at D1-MSN synapses in the SNr with the styryl dye FM1-43 ^29^. This approach has a unique advantage over postsynaptic recordings for the analysis of presynaptic effects of DA and D1R on D1-MSN synapses, since the rate of the fusion-step preceding neurotransmitter release is determined directly and is not obscured by changes in GABA receptor sensitivity in the postsynaptic cell.

FM1-43 was loaded into the lumen of recycling vesicles of D1-MSN synapses by applying a 10Hz stimulus train to the D1-MSN fibers (Fig 1A). After >20 min perfusion of the slice with aCSF containing ADVASEP-7, which removes extrasynaptic FM1-43 label ^30^, FM1-43 fluorescent boutons were imaged by two-photon microscopy. Subsequently, another stimulus train at 2Hz reduced FM1-43 fluorescence with a time-constant of destaining (t_1/2_) corresponding to the rate of synaptic vesicle fusion. In a previous study, we confirmed that this protocol labels D1-MSN boutons ^29^.

**Figure 1.**
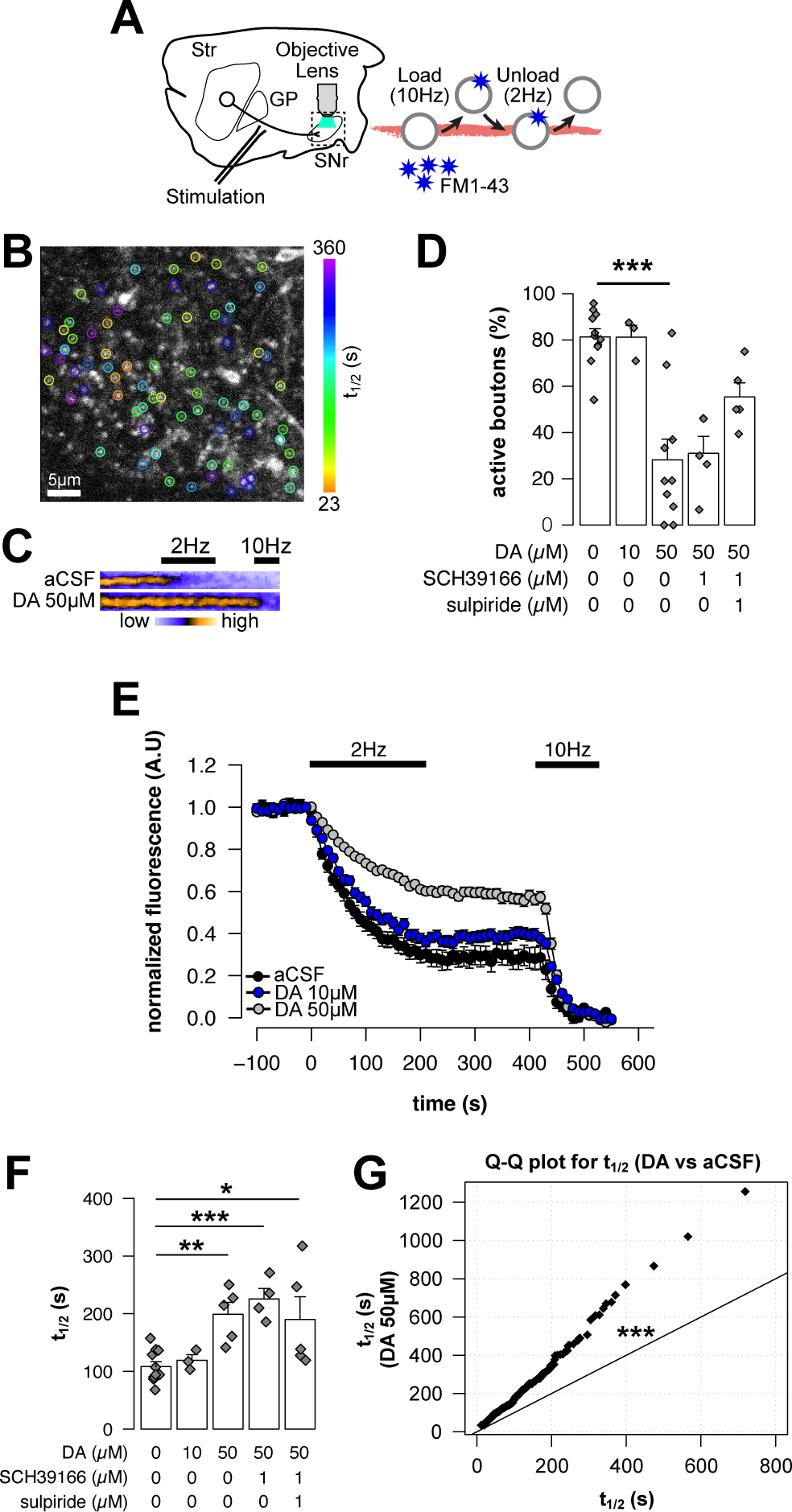
Two-photon imaging of synaptic vesicle fusion in striatonigral synapses reveals inhibition of exocytosis by DA. (A) Schematic illustrating experimental setup employed for measuring the rate of synaptic vesicle fusion at striatonigral boutons using the fluorescent endocytic styryl dye FM1-43. The striatonigral terminals were loaded with FM1-43 via 10Hz electrical stimuli of the striatonigral axons in sagittal mouse brain slices in the presence of D-AP5 (50 µM) and NBQX (10 µM). Subsequently, ADVASEP-7 was applied in the aCSF to remove interstitial FM1-43. The striatonigral axons were then stimulated at 2Hz followed by 10Hz to release dye from the boutons. The fusion of fluorescently labeled synaptic vesicles (FM1-43 destaining) was monitored in real-time using two-photon microscopy. (B) Two-photon image of SNr featuring FM1-43 labeled putative striatonigral boutons. Circles are color-coded to represent the FM1-43 destaining rate (t_1/2_) of each individual bouton. FM1-43 destaining was induced by a series of electrical stimuli applied to the striatonigral axons at a frequency of 2Hz. (C) Two-photon images of FM1-43 labeled striatonigral boutons in an aCSF (top) and DA (bottom) treated brain slice. A decrease in FM1-43 fluorescence was observed in aCSF and DA-treated slices in response to 2 Hz and 10 Hz electrical stimulation, respectively. (D) Percentage of FM1-43 destaining boutons (2Hz vs 10Hz) per slice in aCSF, and in slices treated with DA added to the aCSF alone or 15 min after adding antagonists for D1R (SCH39166) and D2R (sulpiride). Diamonds represent individual slices, while columns show Mean+/-SEM of slices (n = 3-10 slices per condition). There was a significant effect of treatment (Kruskal-Wallis, *χ*^2^(2) = 20.9, p > 0.001). *** p < 0.001 vs. aCSF, Holm’s test. (E) Mean time course of FM1-43 destaining for individual striatonigral boutons in aCSF (black, n = 291 boutons), and aCSF with 10 µM (blue, n = 87 boutons) or 50 µM DA (grey, n = 215 boutons). Mean+/- SEM (3-10 slices per condition). Black bars indicate onset, duration, and frequency of electrical stimulation of the striatonigral axons. (F) The FM1-43 destaining rate (t_1/2_) at striatonigral synapses, during electrical stimulation at 2Hz, examined in various treatment conditions: aCSF alone, aCSF with DA, and aCSF with DA added after >15 min pre-incubation with DA receptor antagonists. Diamonds represent the median t_1/2_ of individual boutons within a slice (range = 6-112 boutons), while columns show Mean+/-SEM of slices (n = 3-11 slices). There was a significant treatment effect (ANOVA, F_(4,23)_ = 7.05, p < 0.001). *p < 0.05, **p < 0.01, ***p < 0.001 vs ACSF, Holm’s test. (G) Quantile-quantile (Q-Q) plot showing the quantiles of the FM1-43 destaining rate, t_1/2_, in slices treated with DA (50 µM) as a function of the quantiles of the FM1-43 destaining rate in aCSF (control) slices. The 45*°* line indicates the relationship between two identical distributions. DA shifts the distribution of the striatonigral boutons to longer t_1/2_, indicating uniform inhibition of exocytosis. (***, KS-test, D = 0.338, p < 0.0001, n = 206 boutons).

As shown in Fig 1B, the rate of vesicle fusion at individual D1-MSN synapses is considerably heterogeneous, with t_1/2_ ranging from 23s to 360s among the boutons within an optical frame. This heterogeneity in neurotransmitter release indicates that D1-MSN synapses have a broad bandwidth of synaptic strengths and multiple levels of plasticity.

### Modulation of exocytosis from D1-MSNs

We first investigated the impact of presynaptic D1Rs on striatonigral release by superfusing the brain slices with DA at a range of concentrations during stimulus-evoked synaptic activity. Stimulation of the striatonigral axons at 2Hz resulted in FM1-43 destaining in 80% of the striatonigral boutons (Fig 1C, D), consistent with our previous report ^29^. In contrast, in the presence of DA (50 µM), few boutons destained with 2Hz stimulation; however, a set of boutons rapidly responded to subsequent 10Hz stimulation (Fig 1C, D). Importantly, aCSF and DA treated slices had similar numbers of FM1-43 destaining boutons (aCSF = 43±8 vs DA = 42±12 boutons, t_(16)_ = 0.08, p > 0.05), indicating that DA selectively reduced synaptic vesicle fusion with 2Hz stimuli. DA also increased the t_1/2_ of FM1-43 destaining to 2Hz stimuli (Fig 1E, F) and shifted the entire population of boutons to slower destaining rates (Fig 1G), indicating a potent and uniform inhibition of release from D1-MSN synapses.

Surprisingly, however, the reduction in active boutons and the increased t_1/2_ of FM1-43 destaining were not prevented by adding D1R (SCH39166, 1 µM) or D2R (sulpiride, 1 µM) antagonists prior to DA in the aCSF (Fig 1D-F). Moreover, the inhibition of synaptic vesicle fusion exerted by DA also occurred in slices from mice with genetic deletion of *Drd1a* (Drd1a-/-), which lack D1Rs (Fig 2A-C). In these mice, DA (50 µM) reduced the number of active boutons (Fig 2B), and DA (10 µM) increased the t_1/2_ of FM1-43 destaining in a majority of the striatonigral synapses (Fig 2C), responses similar to those in wild-type mice. Together, these results demonstrated that DA inhibits the rate of exocytosis at D1-MSNs in the SNr; however, this action is independent of DA receptors.

**Figure 2.**
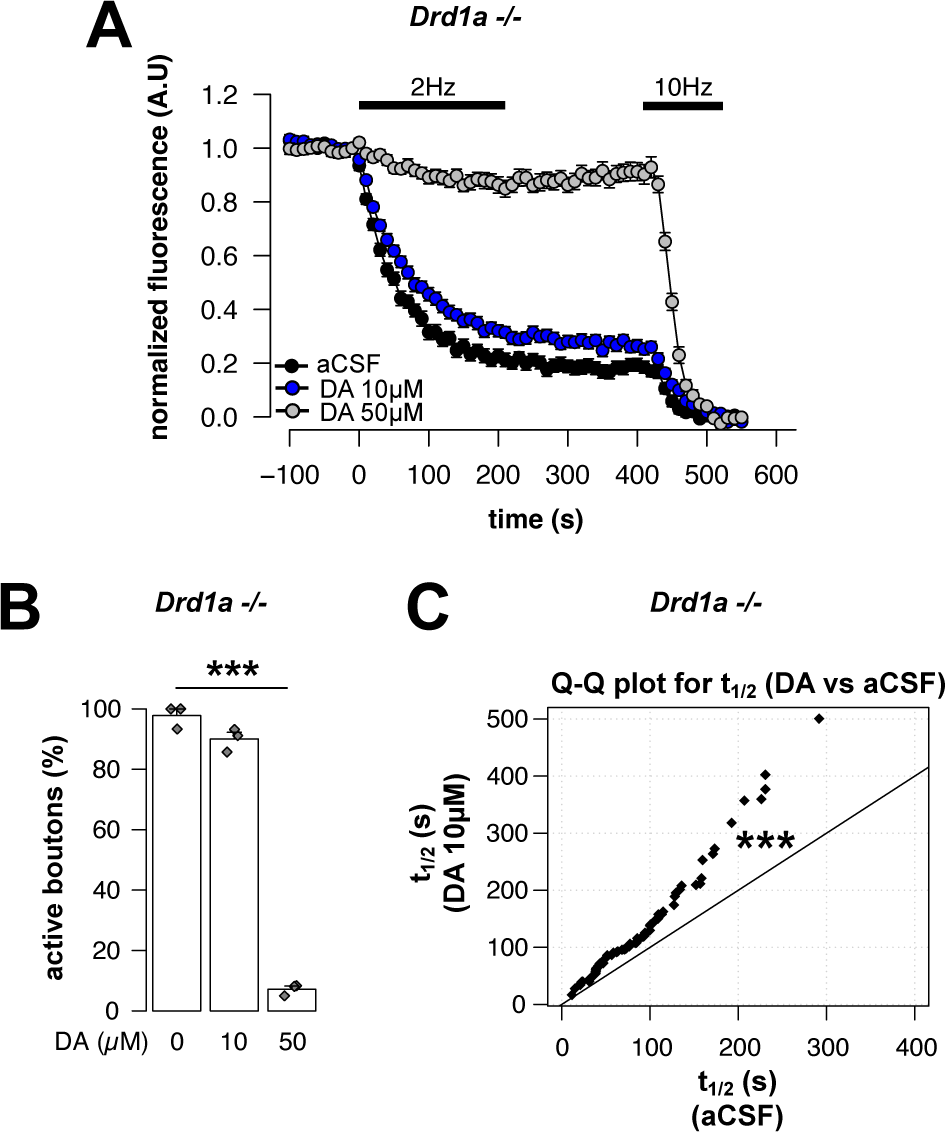
Inhibition of striatonigral exocytosis by DA in *Drd1a*-deficient mice. (A) FM1-43 destaining in slices from *Drd1a*-deficient mice in aCSF (black, n = 81 boutons), aCSF with 10 µM (blue, n = 95 boutons), and 50 µM DA (grey, n = 86 boutons). Mean +/- SEM (n = 3 slices per condition). Black bars indicate onset, duration, and frequency of electrical stimulation of the striatonigral axons. (B) Percentage of FM1-43 destaining boutons (2Hz vs 10Hz) in slices from *Drd1a*-deficient mice treated with aCSF alone or with DA. DA produced a dose-related decrease in 2Hz responsive boutons (Kruskal-Wallis, *χ*^2^(2) = 7.26, p < 0.05, n = 3 slices per condition). ***p < 0.001 vs ACSF, Holm’s test. (C) Quantile-quantile (Q-Q) plot showing the quantiles of the FM1-43 destaining rate, t_1/2_, in slices from *Drd1a*-deficient mice treated with DA (10 µM) as a function of the quantiles of the FM1-43 destaining rate in aCSF (control) slices. The 45*°* line describes the relationship between two identical distributions. Note that DA shifts the distribution of the striatonigral boutons to longer t_1/2_, indicating uniform inhibition of exocytosis. (***, KS-test, D = 0.305, p < 0.001, n = 79 boutons).

To corroborate these findings at pharmacologically relevant concentrations of DA, we added amphetamine (10 µM), which is thought to release DA in the slice through reverse transport via the dopamine transporter, to the aCSF while imaging stimulus-evoked FM1-43 destaining (Fig 3). Like exogenously applied DA, amphetamine reduced the fraction of active boutons (Fig 3A-B) and decreased the rate of the destaining displayed by a significant increase in t_1/2_. Pre-treatment with a D1R antagonist (SKF83566, 1µM) in the aCSF 15 min prior to amphetamine application did not significantly affect the amphetamine-induced decrease in release (Fig 3C).

**Figure 3.**
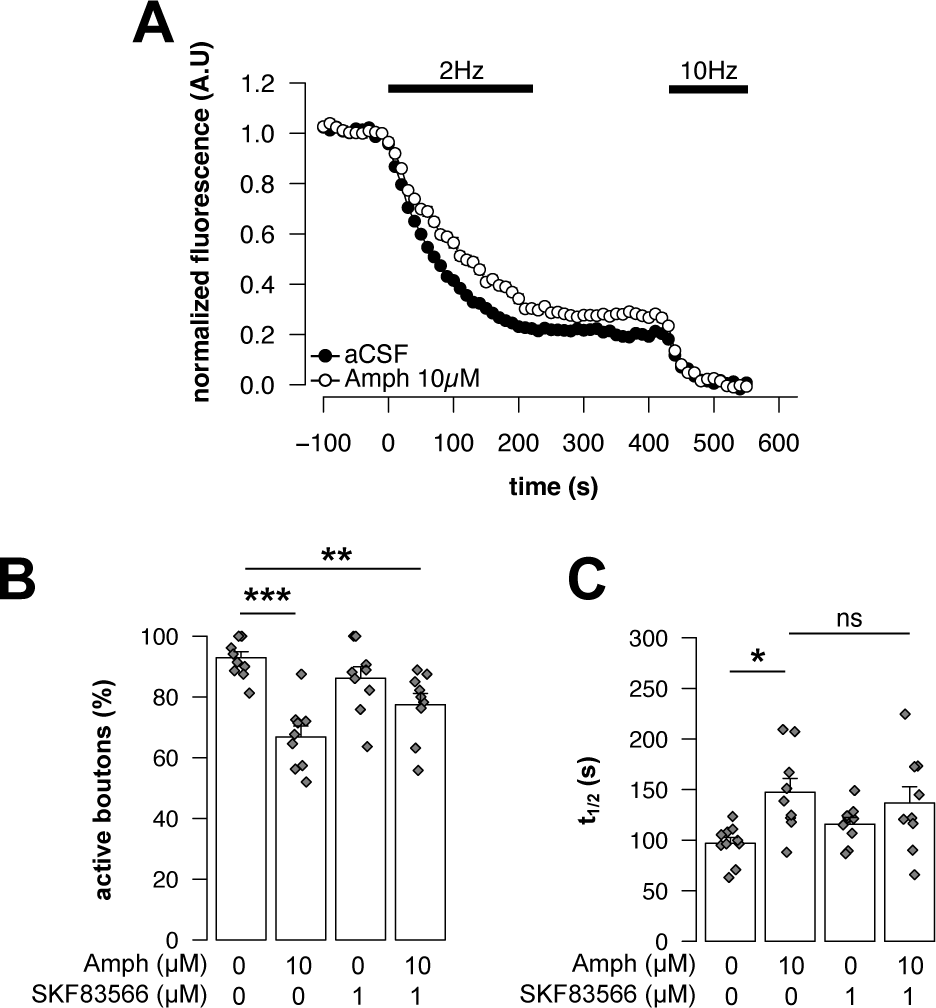
Inhibition of striatonigral exocytosis by amphetamine. (A) FM1-43 destaining in slices in aCSF (black, n = 362 boutons) and aCSF with amphetamine (white, Amph, 10 µM, n = 206 boutons). Mean +/- SEM (n = 9-10 slices per condition). Black bar indicates the onset, duration, and frequency of electrical stimulation of the striatonigral axons. (B) Bar graph shows the ratio of destaining boutons (2Hz vs. 10Hz) per slice in aCSF, and in slices with aCSF and Amph (10 µM), the D1R antagonist (SKF83566, 1µM) or Amph and SKF83566 together. SKF83566 was applied >15 min before Amph. Diamonds represent ratio within individual slices, while columns show Mean+/-SEM of slices (n = 3-10 slices per condition). There was a significant effect of treatment (Kruskal-Wallis, *χ*^2^(3) = 19.56, p > 0.001). ***p < 0.001, **p < 0.01 vs. aCSF, Holm’s test. (C) The FM1-43 destaining rate (t_1/2_) at striatonigral synapses, during electrical stimulation at 2Hz, examined in various treatment conditions: aCSF alone, aCSF with Amph, aCSF with SKF83566, and Amph applied >15 min after SKF83566. Diamonds represent the median t_1/2_ of individual boutons within a slice (range = 201-319 boutons), while columns show Mean+/-SEM of slices (n = 9-10 slices). Two-way ANOVA showed a significant main effect of Amph (F_(1, 33)_ = 10.44, p < 0.001), no main effect of SKF83566 (F_(1, 33)_ = 0.24, p > 0.05) and no Amph:SKF83566 interaction (F_(1, 33)_ = 1.76, p > 0.05), *p < 0.05 vs aCSF, Holm’s test, ns = not significant.

Previous postsynaptic patch-clamp studies in brain slices suggested that D1R agonists increase GABA release in the SNr, based on increased GABA_A_ receptor-mediated synaptic currents in SNr cells ^20, 23, 24^. We therefore examined if selective activation of D1R modulated the rate of synaptic vesicle fusion in the FM1-43-based assay. Two different D1R agonists were tested at a range of concentrations (Fig 4A). Surprisingly, neither the full agonist (SKF81297, 1-10µM) nor the partial agonist (SKF38393, 1-10µM) significantly changed the t_1/2_ of FM1-43 destaining. Moreover, the rate of synaptic vesicle fusion from D1-MSNs was unaffected by D1R blockade (SKF83566, 1µM), adenylyl cyclase activation (NKH477, 10µM), or by DA D2R stimulation (quinpirole, 1µM) (Fig 4B). In summary, our results revealed no evidence supporting a direct effect of D1Rs, or D2Rs, on GABA release from D1-MSNs in the SNr. However, both DA and amphetamine effectively suppressed the rate of synaptic vesicle fusion from D1-MSNs terminals in the SNr.

**Figure 4.**
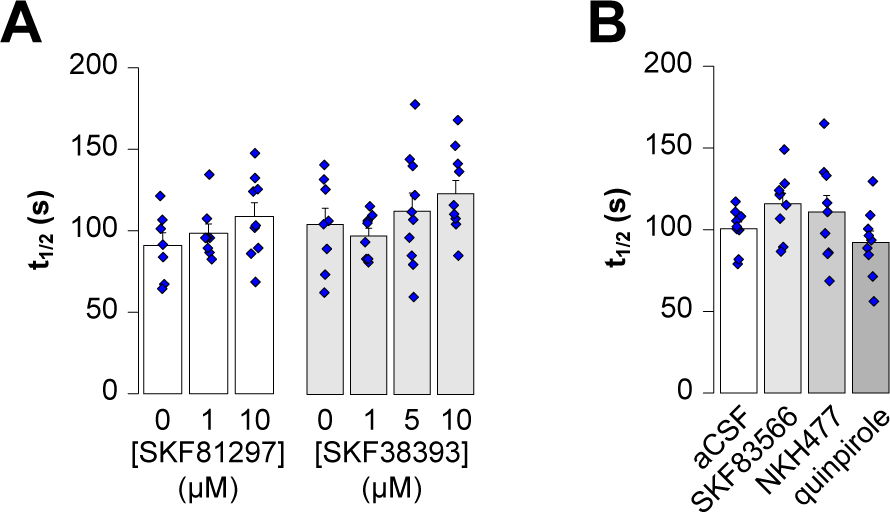
Activation of DA receptor signaling has no effect on striatonigral exocytosis. (A) The FM1-43 destaining rate (t_1/2_) at striatonigral synapses, during electrical stimulation at 2Hz, examined in aCSF, and aCSF with different concentrations of the selective D1R agonists SKF81297 and SKF38393. Diamonds indicate the median t_1/2_ from individual slices, while the columns show Mean+/-SEM of slices (n = 7 – 10 slices per condition). One way ANOVA revealed no significant effect of SKF81297 (F_(2, 21)_ = 1.40, p > 0.05) or SKF38393 (F_(3, 32)_ = 1.50, p > 0.05). (B) The FM1-43 destaining rate (t_1/2_) at striatonigral synapses, during electrical stimulation at 2Hz, examined in different treatment conditions: aCSF, aCSF with D1R antagonist SKF83566 (1 µM), aCSF with adenylyl cyclase activator NKH477 (10 µM) and aCSF with D2R agonist quinpirole (1 µM). Diamonds indicate the median t_1/2_ from individual slices, while the columns show Mean+/-SEM of slices (n = 9 slices per condition). One way ANOVA revealed no significant effect of treatment (F_(3, 32)_ = 2.13, p > 0.05).

### Modulation of Ca2+ influx in D1-MSNs by D1R activation

We have previously reported that FM1-43 destaining depends on Ca^2+^ influx into striatonigral synapses and is prevented by the Ca^2+^ channel blocker Cd^2+^, while the extracellular Ca^2+^ concentration influences the kinetics of FM1-43 destaining ^29^. Thus, the absence of an effect of D1R stimulation on FM1-43 destaining suggests that DA modulates presynaptic Ca^2+^ influx in striatonigral axons through a distinct mechanism.

To investigate this, we conditionally expressed the Ca^2+^ sensitive biosensor, GCaMP, in D1R expressing MSNs by infusing AAVs carrying floxed-GCaMP into the striatum of D1Cre transgenic mice. Histological examination conducted three weeks after surgery confirmed expression of GCaMP in DARPP-32 (a selective MSN marker)-containing striatonigral neurons in the striatum and the axons in the SNr (Fig 5A).

**Figure 5.**
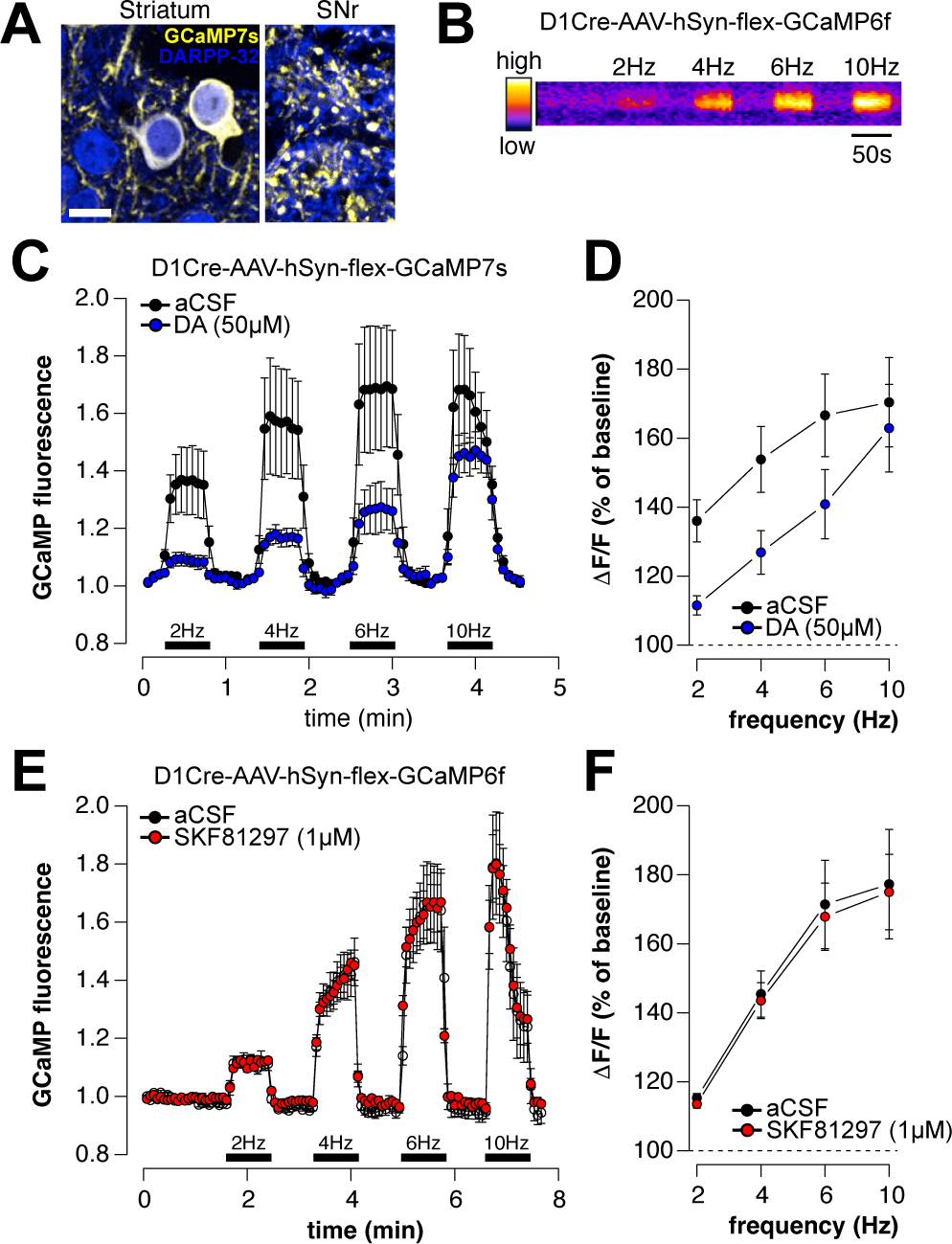
Inhibition of presynaptic Ca^2+^ influx in striatonigral terminals by DA. (A) Confocal images of GFP (yellow) and DARPP-32 (magenta) immunofluorescence in the Striatum (left) and SNr (right) of brain sections from a D1Cre transgenic mouse injected in the striatum with AAV-hSyn-flex-GCaMP7s. Scale bar = 10 µm, 3 µm. (B) Two-photon image of a fluorescent puncta in the SNr of a sagittal brain slice from a *D1Cre* transgenic mouse injected in the striatum with AAV-hSyn-flex-GCaMP6f. Note the increased signal upon electrical stimulation (50 s) of the striatonigral axons at the indicated frequencies. (C) Presynaptic GCaMP7s fluorescence in striatonigral terminals measured using two-photon imaging during electrical stimulation of the striatonigral axons in slices perfused with aCSF (black), and after the application of DA (50 µM, blue). Stimulation-sensitive fluorescent puncta were normalized to their unstimulated fluorescent levels, and the fluorescent values of puncta within slices were averaged. Data shown represents mean+/- SEM of slices (n = 7). Black bars indicate the onset, duration, and frequency of electrical stimulation of the striatonigral axons. (D) Normalized changes in GCaMP fluorescence during electrical stimulation of the striatonigral axons, expressed as a percentage of baseline unstimulated values, in the aCSF (black) and DA-treated slices in C. Two-way repeated measures ANOVA significant main effect of stimulation frequency (F_(4,24)_ = 25.37, p < 0.001), significant main effect of treatment (F_(1,6)_ = 12.4, p < 0.05) and a significant frequency:treatment interaction (F_(4,24)_ = 7.18, p < 0.001). (E) Presynaptic GCaMP7s fluorescence in striatonigral terminals measured using two-photon imaging during electrical stimulation of the striatonigral axons in slices perfused with aCSF (black), and after the application the D1R agonist SKF81297 (1 µM, blue). Stimulation-sensitive fluorescent puncta were normalized to their unstimulated fluorescent levels, and the fluorescent values of puncta within slices were averaged. Data shown represents mean+/- SEM of slices (n = 4). Black bars indicate the onset, duration, and frequency of electrical stimulation of the striatonigral axons. (F) Normalized changes in GCaMP fluorescence during electrical stimulation of the striatonigral axons, expressed as a percentage of baseline unstimulated values, in the aCSF (black) and SKF81297-treated slices in E. Two-way repeated measures ANOVA significant main effect of stimulation frequency (F_(4,12)_ = 22.46, p < 0.001), no significant main effect of treatment (F_(1,3)_ = 0.39, p < 0.58), no significant frequency:treatment interaction (F_(4,12)_ = 0.42, p < 0.79).

As fluorescence in individual presynaptic puncta may be influenced by the transduction efficacy of GCaMP, we reasoned that a suitable approach to quantify the effect of D1R stimulation on Ca^2+^ influx would be to measure the relative increase in fluorescence in response to a gradual increase in stimulation frequency under both control and D1R-stimulated conditions. To do so, we imaged Ca^2+^ transients in the SNr of sagittal brain slices using two-photon microscopy, while applying electrical stimulation to the striatonigral fibers at a range of frequencies.

As shown in Fig 5B, the intensity of GCaMP signal paralleled the frequency of stimulation. Application of DA (50 µM) produced a notably blunted stimulus-related change in GCaMP fluorescence, indicating a potent inhibition of presynaptic Ca^2+^ influx (Fig 5C). In both aCSF control slices and DA-treated slices stimulation with increasing frequency resulted in a gradual increase in the relative GCaMP fluorescence (ΔF/F). However, only at 10Hz, the highest frequency tested, did we observe a significant increase in GCaMP fluorescence compared to baseline levels in DA-treated slices. Thus, the inhibition of presynaptic Ca^2+^ influx by DA, like inhibition of FM1-43 destaining, is frequency-dependent and can be overcome when striatonigral synapses are activated at 10 Hz.

In contrast to DA, application of a D1R agonist (SKF81297, 1µM) had no discernible effect on GCaMP fluorescence compared to aCSF alone (Fig 5E). The lack of effect of the D1Rs was evident in the relative GCaMP fluorescence (ΔF/F) over a range of levels of increased synaptic activation (Fig 5F).

In summary, our presynaptic GCaMP recordings revealed that striatonigral synapses were markedly inhibited by DA but were unaffected by D1R activation, consistent with the effects observed in the synaptic vesicle fusion measurements above.

### Postsynaptic recordings in SNr cells

Because our results contrast with those from prior postsynaptic recordings ^20, 23, 24^, we then investigated the effect of D1Rs in the SNr by performing patch-clamp recordings on SNr cells. To achieve specific activation of striatonigral synapses, we transduced the light-activated cation channel channelrhodopsin (ChR2) in D1-MSNs by infusing an AAV-DIO-ChR2 into the striatum of D1Cre transgenic mice (Fig 6A). Subsequently, we prepared sagittal brain slices and recorded SNr neurons in voltage-clamp configuration, using recording pipettes containing 144 mM chloride to enhance the detection of inward inhibitory postsynaptic currents. Prior to whole-cell recordings, we measured the tonic firing activity in cell-attached mode and analyzed only neurons with the characteristic firing rate of SNr GABA neurons ^15^ for further analysis.

**Figure 6.**
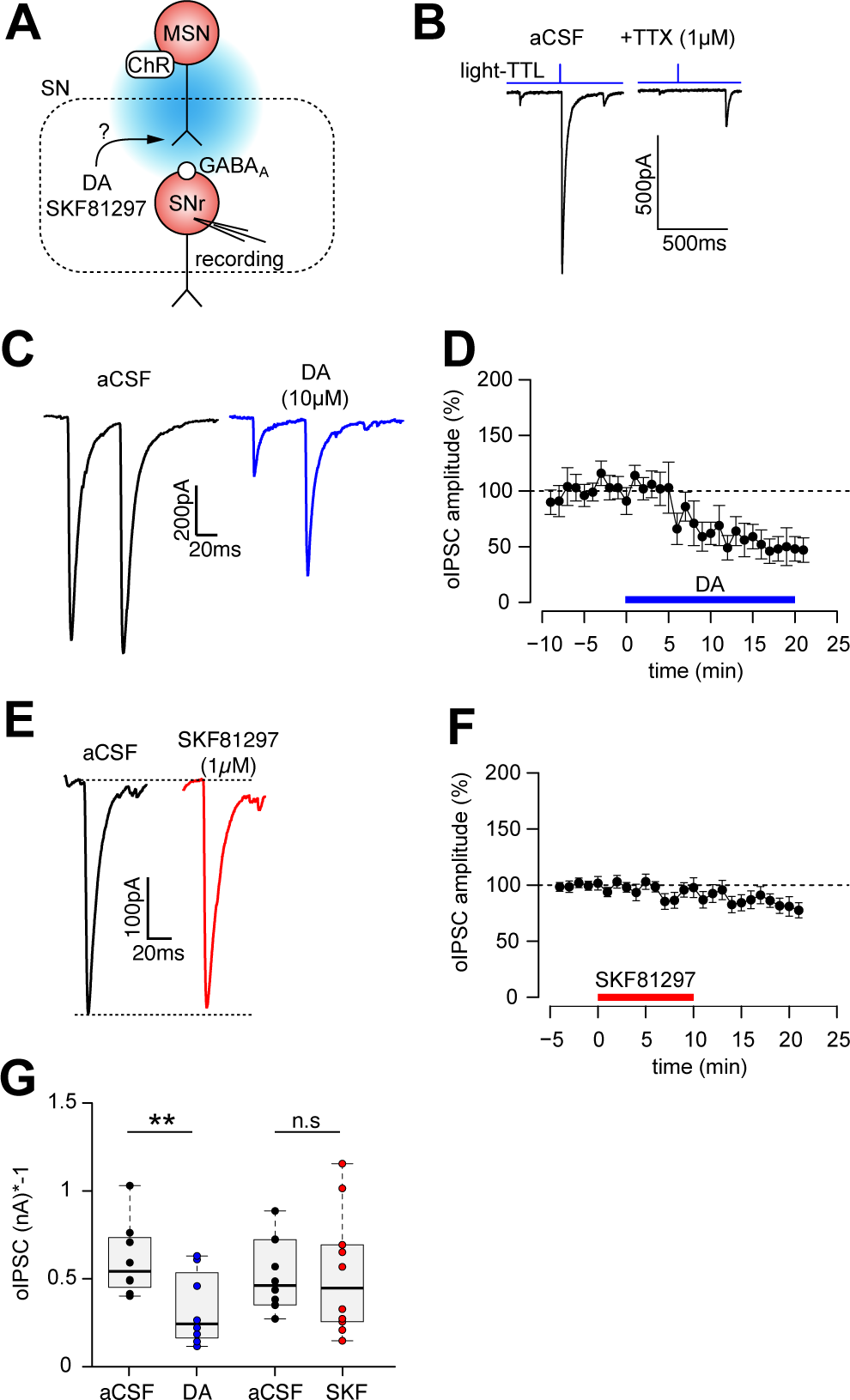
Inhibition of striatonigral GABA release by DA. (A) Schematic depicting the experimental setup employed to examine the effect of DA receptor stimulation on striatonigral GABA release. D1Cre transgenic mice received intrastriatal injections of AAV-hSyn-DIO-hChR2(H134R) to express the light-activated cation ChR in striatonigral efferents. Optogenetically induced GABA release was recoded indirectly by measuring GABA_A_ receptor currents in neurons of the SNr in whole-cell patch clamp configuration. DA receptor ligands were introduced into the aCSF. (B) Current responses measured in an SNr neuron following a blue flash delivered to the slice, in aCSF (left) and in aCSF containing the sodium-channel blocker tetrodotoxin (TTX, 1µM, right). The timing of the digital TTL step used to deliver the blue flash is indicated (light-TTL). TTX effectively blocked the release of GABA and eliminated the synaptic current. (C) Recorded current responses from an SNr neuron in a brain slice subjected to a pair of blue flashes (100 µs) delivered with 50 ms interstimulus interval in aCSF (black) and in aCSF with DA (10 µM, blue). (D) Inhibitory postsynaptic current amplitude (oIPSC) elicited by intermittent photo-stimulation of striatonigral synapses. The aCSF contained DA (10 µM) at the indicated interval (blue). Currents were normalized to the pre-DA application average. Data represents the mean+/-SEM in percent, derived from recordings of n = 9 SNr neurons. (E) Recorded current responses from an SNr neuron in a brain slice subjected to a blue flash (100 µs) in aCSF (black) and in aCSF with the D1R agonist SKF81297 (1 µM, red). (F) Inhibitory postsynaptic current amplitude (oIPSC) elicited by intermittent photo-stimulation of striatonigral synapses. The aCSF contained the D1R agonist SKF81297 (1 µM) at the indicated interval (red). Currents were normalized to the pre-SKF81297 application average. Data represents the mean+/-SEM in percent, derived from recordings of n = 10 SNr neurons. (G) Box and whisker plot showing the average inhibitory postsynaptic current (oIPSC) elicited by photo-stimulation of striatonigral synapses in different treatment conditions. In DA containing aCSF, photo-stimulation produced significantly smaller oIPSCs than in aCSF (t_(7)_ = 4.24, p < 0.01, paired *t-*test). oIPSCs recorded in aCSF with SKF81297 did not differ from aCSF alone (t(9) = 0.13, p > 0.05, paired T-test). Symbols represent oIPSC in individual cells. ** p < 0.01, n.s = not significant.

Brief photo-stimulation (0.1 – 0.5 ms) of the striatonigral fibers, approximately 1 mm rostral to the SNr, elicited optical inhibitory postsynaptic currents (oIPSC) in SNr neurons. These oIPSCs were completely blocked by TTX (1 µM, Fig 6B) and picrotoxin (100 µM, not shown), confirming that they consisted of a GABA_A_ current produced by the depolarization-induced GABA release from D1-MSN axons.

After establishing a stable current response in aCSF, we applied DA (10 µM) and subjected the slice to intermittent (0.05 Hz) photostimulaton. Within 5 min, DA robustly decreased the oIPSC by approximately 50% (Fig 6 C-D). The response persisted for as long as DA was present in the aCSF. Subjecting the slices to the D1R agonist SKF81297 (1 µM) in aCSF did not change the amplitude of the striatonigral oIPSC in SNr cells (Fig 6 E-F). In contrast to SKF81297, DA produced a significant reduction in the oIPSC amplitude compared with drug-free aCSF (Fig 6G). These results are consistent with the findings obtained with presynaptic Ca^2+^ and synaptic vesicle fusion optical recordings and confirm that DA but not D1R activation modulates GABA release from striatonigral synapses.

### DA inhibits striatonigral release through 5-HT1B receptors

Also consistent with the results of FM1-43 destaining, we observed that the inhibitory effect of DA on the striatonigral oIPSC persisted in slices pre-incubated with DA receptor antagonists (SCH23390 1 µM and sulpiride 1 µM, not shown). A possible explanation this action is that DA influences other neurotransmitters that inhibit striatonigral GABA release. In SNr, DA enhances 5-HT neurotransmission ^31^, which in turn reduces striatonigral GABA release through the activation of 5-HT1B receptors ^32, 33^.

To investigate if DA depresses striatonigral GABA release through a mechanism involving 5-HT1B receptors, we recorded the modulation of oIPSCs by DA in SNr neurons in the presence of the 5-HT1B antagonist NAS-181 (Fig 7A). Application of NAS-181 in the aCSF (5 µM) produced a non-significant effect on striatonigral oIPSCs relative to aCSF alone (Fig 7B-D). Subsequent addition of DA (10 µM) did not further decrease the striatonigral GABA release (Fig 7B-D). In conclusion, the potent inhibitory effect of DA on the striatonigral oIPSC was completely blocked by NAS-181. Altogether, these results suggest that the ability of DA to decrease striatonigral GABA release depends on activation of presynaptic 5-HT1B receptors on striatonigral terminals.

**Figure 7.**
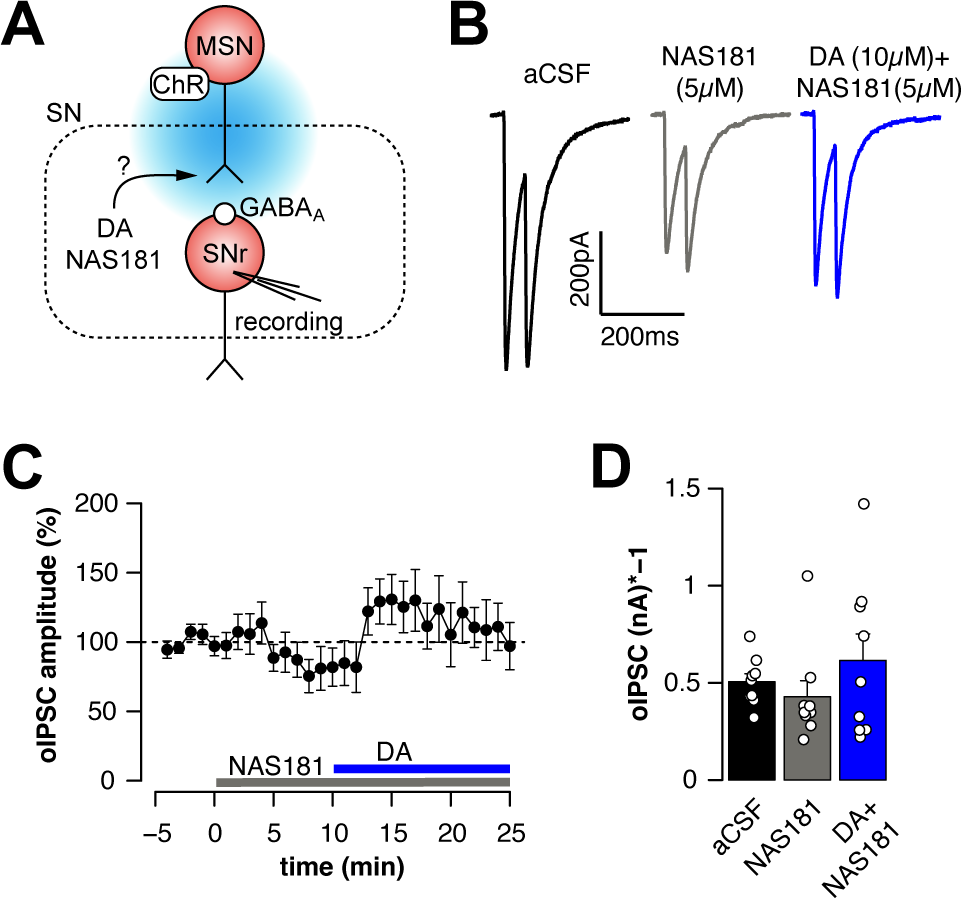
Antagonism of 5-HT1B receptors prevents inhibition of striatonigral GABA release by DA. (A) Schematic depicting the experimental setup employed to examine the effect of NAS-181 on striatonigral GABA release in brain slices from mice expressing ChR2 in D1-MSNs. (B) Current responses in a voltage-clamped SNr neuron to a pair of photostimuli (100 µs) applied to striatonigral synapses at 50 ms interstimulus interval in different treatment conditions: in the aCSF (black), aCSF with the 5-HT1B receptor antagonist NAS-181 (5 µm, grey), and aCSF with NAS-181 together with DA (10 µm, blue). (C) Inhibitory postsynaptic current response (oIPSC) elicited by intermittent photo-stimulation (100 µs) of striatonigral synapses. The aCSF contained NAS-181 (5 µM, grey) and NAS-181 together with DA (10 µM, blue) at the indicated intervals. Currents were normalized to the pre-NAS-181 application average. Data represents the mean+/-SEM in percent, derived from recordings of n = 10 SNr neurons. (D) Bar plot showing the average current amplitude in voltage-clamped SNr neurons following photo-stimulation of striatonigral synapses in aCSF (black), in aCSF with NAS-181(grey), and in aCSF with NAS-181 together with DA. Symbols represent the average current before and 10 min after application of NAS-181 alone, and 10 min in with NAS-181 together with DA in individual SNr neurons. Columns show means +/- SEM of neurons (n = 6). One-way repeated measures ANOVA found no significant effect of treatment (F_(2, 10)_ = 0.83, p > 0.05).

## Discussion

We report that both DA and amphetamine decrease evoked GABA release from striatonigral axons. Surprisingly, the activation of DA receptors with synthetic D1R and D2R agonists had negligible effects on striatonigral GABA release, as did chemical activation of adenylyl cyclase – a crucial component of the canonical signaling cascade utilized by DA receptors. The inhibitory effect of DA persisted in the presence of DA receptor antagonists and was intact in D1R-deficient mice, indicating a non-DA receptor action. Instead, we found that the effect of DA was prevented by inhibiting 5-HT1B receptors, consistent with a previous suggestion that DA decreases GABA release in the striatum by activating 5-HT1B receptors ^28^. Together, our study indicates that 5-HT1B receptors mediate inhibitory DAergic actions on GABA release from MSN axons in the striatum and the midbrain, two regions involved in the DAergic control of volitional movement orchestrated by the basal ganglia.

### Inhibition of striatonigral GABA release by DA depends on regulation of presynaptic Ca^2+^ influx and 5-HT1B actions

Our study revealed an inhibitory effect of DA on three crucial steps of the activity-dependent GABA release from striatonigral axons: a decreased Ca^2+^ influx into striatonigral axons during synaptic activation observed through two-photon imaging of axonal GCaMP in D1-MSNs; a dose-dependent reduction in synaptic vesicle fusion quantified in two-photon images of FM1-43 destaining from striatonigral boutons; and decreased amplitude of GABA_A_ receptor currents in SNr GABA neurons following selective optogenetic activation of D1-MSN axons. Collectively, these findings provide independent evidence for the inhibitory action of DA on striatonigral GABA release.

These findings are in apparent disagreement with earlier studies ^20, 34^, which showed that DA increases GABA_A_ synaptic currents produced by stimulation of the SNr, and enhances depolarization-induced release of radiolabeled GABA from midbrain slices. These studies attributed the effect of DA to D1R activation ^20, 34^. In contrast, our investigation focused on the presynaptic effects of DA on the striatonigral axons, and are consistent with a previous report indicating that DA reduces GABA_A_ currents recorded in SNr GABA neurons elicited by electrical stimulation of SNr afferents ^22^.

We were surprised to find that antagonists targeting D1R and D2R-like receptors failed to prevent the presynaptic inhibitory action of DA. Instead, we demonstrate that the presynaptic inhibition of GABA release by DA requires activation of 5-HT1B receptors, which are highly expressed on striatonigral terminals^35^. 5-HT1B receptors have been shown to mediate inhibition of striatonigral GABA release produced by serotonin ^32, 33^. In addition, these receptors decrease GABAergic synaptic transmission between MSNs in the ventral striatum induced by DA ^28^ and inhibit GABA release from D2-MSN axons in the ventral pallidum induced by cocaine, a psychostimulant that increases extracellular DA ^36^.

Our observation is consistent with a growing body of evidence demonstrating interactions between DAergic and serotonergic neuromodulation in the central nervous system ^37, 38^. Such interplay could be particularly relevant in the SNr where a dense serotonergic innervation from the dorsal raphe nucleus ^39–41^ influences inhibition of striatonigral GABA release via 5-HT1B receptors and excitation of SNr GABA cells through 5-HT2C receptors ^33, 42^.

The specific mechanism by which DA inhibits striatonigral release of GABA via 5-HT1B receptors is unclear. One possibility is that DA directly binds to 5-HT1B receptors, which is supported by the capacity of DA, along with synthetic ligands for DA receptors, to bind to several 5-HT receptor isoforms ^43^. Alternatively, the effect of DA could be mediated by increased serotonin release ^31^, which would then activate 5-HT1B receptors on striatonigral terminals, subsequently decreasing GABA release. Another possible indirect action of DA is suggested by the observation that DA can enter serotonin terminals through the serotonin transporter ^44^ and promote serotonin release ^45^, presumably by displacing serotonin from VMAT2 containing synaptic vesicles ^46^. To our knowledge, whether this mechanism is unique to SNr slices exposed to exogenous DA ^31^ is yet to be evaluated.

An involvement of 5-HT1B receptors in the effect of DA is further substantiated by our observation that inhibition of Ca^2+^ influx and the FM1-43 destaining are frequency dependent, with the effect of DA on Ca^2+^ influx decreased as the electrical stimulation was increased from 2Hz to 10Hz. Similarly, FM1-43-labelled vesicles that remained with 2Hz electrical stimulation rapidly disappeared after 10Hz stimulation. These DA-dependent effects resemble those of baclofen, a GABA_B_ receptor agonist that inhibits FM1-43 destaining at striatonigral ^29^ and hippocampal synapses ^47^. The frequency-dependent inhibition by Gi protein-coupled receptors, such as 5-HT1B and GABA_B_ receptors, is attributed to the dissociation of the G*βγ* subunits from its binding to voltage gated Ca^2+^ channels ^47^, enabling Ca^2+^-mediated influx and GABA release during high levels of synaptic activity. These results further support that DA modulates striatonigral activity through presynaptic Gi protein-coupled 5-HT1B receptors in the SNr, in contrast to G_olf_ protein-coupled D1R receptors.

### Possible functions of D1R on striatonigral terminals

Most D1Rs in the SNr originate from the striatum ^48^ and are positioned extrasynaptically on striatonigral efferents ^7, 9^. Considering the abundance of D1R expressed on striatonigral axons, it is reasonable to hypothesize that D1Rs influence striatonigral GABA release. Our presynaptic methods revealed, however, that the activation of D1Rs with synthetic agonist does not modify the activity of striatonigral synapses. Similarly, the postsynaptic GABA_A_ currents recorded in SNr GABA neurons after selective optogenetic activation of striatonigral synapses were not affected by D1R activation. These findings are surprising in light of previous studies showing that D1R agonists increase SNr GABA neurotransmission ^18–20, 23, 24, 34, 49, 50^.

Since previous studies investigated GABA release indirectly, it remains possible that the detected GABA originates from multiple sources that are differentially sensitive to D1R stimulation. For example, upon SNr activation by either depolarizing solution ^34, 50^ or electrical stimulation ^20^, the SNr GABA neurons, which express D1Rs ^51^ could release GABA through axon collaterals ^52–54^. The possibility that SNr GABA neurotransmission is enhanced by activation of postsynaptic D1Rs is supported by reports showing that D1R activation depolarizes SNr GABA neurons ^21,51^. Other potential sources of GABA include dendritic GABA release, a well-studied feature of thalamic interneurons ^55^, and GABA release from SNr astrocytes ^49^, which also express D1R ^56^. SNr stimulation also activates pallidonigral synapses ^25, 57^, which are inhibited by D2R activation ^23^. In our study, quinpirole, a D2R agonist, had no effect on FM1-43 destaining and we conditionally expressed GCaMP and ChR2 in D1-MSNs, indicating that pallidonigral synapses contributed minimally to our presynaptic measurements ^29^.

We note that the conclusion that D1Rs do not modulate GABA release from striatonigral synapses is valid for the synchronous activation of synaptic release, driven by action potentials and the opening of voltage gated Ca^2+^ channels. If the modulation of GABA release by D1Rs is not reliant on Ca^2+^, it would escape detection in these experiments. This possibility is not ruled out under some conditions, as DA and D1R agonists increase SNr GABA release in the presence of TTX and Ca^2+^ channel blockers ^20^, which indicates that D1R stimulation increases asynchronous GABA release. The enhanced GABA release attributed to D1Rs could result from their ability to activate cAMP signaling. Enhanced cAMP signaling, either through forskolin-activated adenylyl cyclase ^20, 58^ or by inhibition of phosphodiesterase activity ^59^ potentiates GABA currents in SNr cells, thus facilitating GABA neurotransmission. Increased cAMP signaling triggered by D1R activation could influence striatonigral synaptic function in multiple ways, including via vesicular recycling ^60^, regulation of the size of the recycling pool of vesicles ^61^, and facilitating the recruitment of silent synapses ^62^ - mechanisms of synaptic facilitation independent of instantaneous presynaptic Ca^2+^ influx ^63^. If these changes occur slowly, they may not be measurable in the acute brain slice preparation.

The complex regulation of SNr GABA release from different sources motivates further studies using methods that allow for a systematic analysis of the target of facilitatory D1R effects.

### Functional implications for motor control and movement disorders

DA released from nigrostriatal synapses regulates basal ganglia output through dichotomous modulation of D1-MSNs and D2-MSNs ^1^. According to the to the rate model of the basal ganglia-mediated control of movement, this promotes motor activity upon inhibition of SNr neurons ^64, 65^. Here, our evidence supports the idea that DA influences behavior by regulating D1-MSN activity locally within the SNr ^4, 17, 66^. Rather than controlling the rate of SNr firing, we propose that DA released from SNc dendrites into the SNr participates in activity-based selection of striatonigral terminals, determining the pattern of SNr firing involved in action selection by the basal ganglia ^67^.

SNr GABA neurons inhibit SNc DA neurons via collaterals of the axons projecting towards motor centers in the thalamus and brain stem ^52, 68–70^. Disinhibition of SNc DA neurons following striatonigral-mediated reduction of SNr firing causes DA neurons to burst ^71, 72^, promoting DA release in the striatum ^73, 74^, which facilitates firing of MSNs upon cortical activation *in vivo* ^75^. Our results suggest that DA in SNr increases GABA release during MSN burst firing due to short-term facilitation of striatonigral synapses ^26^. The frequency-dependent regulation of GABA release in the SNr by DA presents a novel mechanism through which DA can facilitate salient motor behaviors within the basal ganglia functional loops ^76^.

The midbrain circuit, comprising the DA neurons of the SNc and GABA neurons of SNr alongside high levels of both DA and serotonin receptors, could also play crucial roles in the emergence of motor symptoms of Parkinson’s disease and their response to levodopa - the mainstay pharmacotherapy for Parkinsonian symptoms. For example, in a mouse model with progressive DA pathology, parkinsonian disabilities such as akinesia and gait disturbances appear after the reduction of DA release in the SNr ^77^. Loss of DA in the SNr increases the probability of striatonigral GABA release, which drives sensitized motor responses in parkinsonian mice after optogenetic activation of D1-MSN terminals in the SNr ^29^. Based on our findings, this increased striatonigral GABA release in DA depleted animals can be explained as a loss of DA-mediated frequency-dependent filtering of striatonigral activity and may explain how levodopa triggers sensitized motor responses in Parkinson’s disease, such as dyskinesia, dopamine dysregulation syndrome or impulse disorders. Emerging evidence indicates that dyskinesia is caused by dysregulated DA release from serotonin terminals ^38^, and is further substantiated by the antidyskinetic effects of 5-HT1 receptor agonists ^78, 79^. Our findings show that such effects could be mediated through DA interactions with serotonin receptors in the SNr, suggesting a mechanism for further development of antidyskinetic therapeutic strategies.

## Methods

### Experimental subjects

Experiments were performed on mice of both sexes, aged 5 weeks and older. Wild-type C57Bl/6J mice were purchased from Jackson Laboratories (Bar Harbor, ME, USA). Tg(Drd1a-cre)EY262Gsat/Mmcd were from the GENSAT project at Rockefeller University ^80^. Heterozygous Drd1a wt/ko breeders ^81^ were a generous gift from the late Dr. Marc G. Caron (Duke University). Transgenic mice were bred into the C57Bl/6J background for more than 10 generations and experimental subjects were the offspring heterozygous breeding pairs. All animal experiments approved by the Institutional Animal Care and Use Committee (IACUC) at the Columbia University and the Swedish Board of Agriculture (License: 18194-2018) and followed the guidelines of the Research Ethics Committee of Karolinska Institutet, Swedish Animal Welfare Agency and the European Communities Council Directive 2010/63/EU.

### Surgical procedures

Mice were anesthetized with isoflurane 4% and positioned in a stereotaxic frame (Stoelting) equipped with a thermal pad to maintain normothermia. Anesthesia was maintained with 1-2% isoflurane. All animals were injected subcutaneously with 0.1 mg/kg Temgesic at the start of the surgery. For the stereotaxic virus vector injections, 250 nl of AAV5-DIO-ChR2-mCherry, provided by Karl Deisseroth (Addgene, 20297) and AAV5-Syn-Flex-GCaMP6f or AAV-Syn-FLEX-jGCaMP7s, provided by Douglas Kim (Addgene, 100833, 104491), was infused with a heat-pulled glass micropipette attached to an injection apparatus (Stoelting). The injection was targeted at the striatum using the following coordinates relative to bregma and the dural surface: anteroposterior +0.8 mm, mediolateral -1.7 mm and dorsoventral - 2.1 mm. Following the AAV infusion, the micropipette was left in place at the injection site for at least 5 min to allow diffusion of the viral particles and prevent backflow along the micropipette tract. Finally, the incision was closed with self-absorbed sutures (Vicryl Rapid, 5-0, FS-2) and the mice were allowed to recover from anesthesia on a thermal pad. The mice received postoperative pain relief with bidaily injections with Rimadyl (5 mg/kg s.c) for a period of 2 days following the procedure.

### Brain slice preparation

Mice were sacrificed by cervical dislocation and sagittal slices (250-400 *μ*m) were prepared with an angle of 10-15*°* to maintain rostrocaudal striatonigral projections ^82^ using a vibrating microtome (Leica VT1200). Slice preparation was performed in ice-cold sucrose-based artificial cerebrospinal fluid (ASCF) containing (in mM): 180 sucrose, 10 NaCl, 2.5 KCl, 25 NaHCO_3_, 0.5 CaCl_2_, 7 MgCl_2_, 1.25 NaH_2_PO_4_, 10 glucose (pH 7.3, 300-305 mOsm/l). Following preparation, the slices were left to equilibrate for 30 min in 34*°*C aCSF containing (in mM): 125 NaCl, 2.5 KCl, 25 NaHCO_3_, 2 CaCl_2_, 1 MgCl_2_, 1.25 NaH_2_PO_4_, 10 glucose (pH 7.3, 300-305 mOsm/l), and were then transferred to room temperature (21-23°C) where they were kept until experimentation. In the two-photon imaging experiments with GCaMP7 and patch-clamp recordings, we prepared slices according to the N-Methyl-D-glucamine (NMDG) protective recovery method ^83^. The slices were submerged in a recording chamber and continuously perfused at rate of 1.5 – 2 ml/min in aCSF. A slice anchor was placed on top of the slice to prevent drift. All solutions were saturated with carbogen-gas (95% O_2_ and 5% CO_2_) during slice preparation and experiments.

### Patch clamp electrophysiology

Recordings were performed on a Slicescope Pro 2000 electrophysiology system (Scientifica) equipped with a MultiClamp 700B amplifier and digidata 1550B analog-to-digital converter connected to a personal computer running pClamp 11.0.3 (Molecular Devices). Brain slices were illuminated with a 780 nm LED module. Cells were visualized with a 40 × 0.8 NA water-immersion objective (LUMPlanFLN, Olympus) and Dodt gradient contrast tube optics, and projected to a personal computer with an infrared-sensitive CCD SciCam Pro camera. Recording pipettes were fabricated with a vertical puller (Narishige PC-100) and had a resistance of 3-5 MOhm when filled with internal solution composed of (in mM): 140 CsCl, 2 MgCl2, 10 HEPES, 2 EGTA, 2 MgATP and 3 QX-314. pH was adjusted to 7.2 with CsOH. Recordings were performed at 32 °C in aCSF. Neurons were clamped at -70 mV (Liquid junction potential not corrected) and analysis was restricted to cells with stable (< 10% change) access resistance below 20 MOhm, as assessed by a hyperpolarizing (-10 mV) step command in each sweep. Photosensitive synaptic currents were evoked in brain slices from D1-Cre-AAV5-ChR2 mice by delivering flashes (0.1 – 0.5 ms) of blue light from a CoolLED PE-300 Ultra LED module through the 40x objective.

### Two-photon laser scanning microscopy

FM1-43 and GCaMP (ver 6f and 7s) imaging were performed on a Prairie Ultima (Prairie Technologies, Middleton, WI, USA) and a Scientifica Hyperscope (Scientifica, Uckfield, UK) multiphoton imaging system, respectively. Two-photon excitation was provided by a tunable Chameleon Vision 1 femtosecond Ti-Sapphire laser (Coherent) and the fluorescence passed through T565lpxr, ET525/50m and ET620/60m filters (Chroma Technologies) before being measured by a set of bi-alkali photomultiplier tubes (PMT, R9880U, Hamahatsu). The laser was tuned to 900 nm and 940 nm to excite FM1-43 and GCaMP, respectively. The laser power delivered to the sample was adjusted by a voltage-controlled pockel cell and was maintained below 15 mW at the sample to minimize phototoxicity and photo bleaching. To activate striatonigral axons, a concentric bipolar electrode was positioned in the slice, a few µm into the cerebral peduncle at approximately 1 mm rostral to the SNr. Stimulus length (200 µs) and frequency were controlled by an Arduino Uno connected to two ISO-Flex (AMPI, Israel) stimulus isolators to deliver intermittent bipolar current pulses of ± 300 µA.

#### FM1-43 loading, destaining and analysis

FM1-43 imaging of vesicular fusion at striatonigral synapses was conducted following a previously outlined protocol ^29^. In brief, FM1-43, dissolved in aCSF (10 µM), was applied to an acute brain slice for 15 minutes in the presence of NBQX (10 µM) and AP-5 (50 µM). Incorporation of FM1-43 into recycling vesicles was done by electrically stimulating the striatonigral fibers at 10 Hz for 5 min. After at least 20 minutes of washout with aCSF containing 100 µM Advasep-7 (Biotium), a cyclodextrin that minimizes extraneous FM1-43 labeling ^30^, the slice was subjected to 2Hz electrical stimulation for 4 min to destain and unload FM1-43. Finally, to ensure complete depletion of the FM1-43 labeled vesicle pool, the slice was then subjected to 10Hz stimulation for 3 minutes.

We acquired Z-stacks consisting of 5 images (16 bit, 333 x 333 pixels) separated by 1 µm at sampling frequency 0.1Hz using PrairieView 4.0.50. The field of view was 37.5 x 37.5 µm (pixel size = 0.113 µm) and dwell time 5.2 µs/pixel, and images were aquired at 40-80 µm into the slice. Images were imported into Fiji ^84^ as time-series and inspected for displacement in the x-y-z axes. Only time-series exhibiting minimal slice movement were selected for further analysis using Fiji and R version 4.1.2 ^85^. The z-stacks were collapsed as a maximum intensity projection, and then registered with the StackReg plugin ^86^. Putative synaptic boutons were selected with the analyze particles feature. Regions of interest (ROIs) were delineated around fluorescence puncta using predetermined parameters: 1) 1.5 times the standard deviation of the fluorescence intensity of the background, assigned from an area in the image lacking evident fluorescence; 2) circular shape; and 3) diameter ranging from 0.1 to 1.5 µm. The mean fluorescence intensity was extracted for each ROI across the time-series. Fluorescence run-down during imaging (due to ADVASEP-7) was corrected for each time point by dividing the mean fluorescence intensity of the ROI with the corresponding fluorescence value of the area outside the ROIs ^87^. The ROI values were then normalized to the fluorescent intensity just before electrical stimulation (t = 0; F_0_) and after the electrical stimulation with 10Hz (F_inf_). The criteria differentiating active synaptic boutons from those showing insignificant FM1-43 destaining were determined empirically by measuring the minimal FM1-43 destaining in aCSF with Cd^2+^ (200 µM), an inhibitor of exocytosis dependent on presynaptic Ca^2+^ influx, and in slices exposed to Ca^2+^ free aCSF ^29^. ROIs exhibiting >10% reduction in raw FM1-43 fluorescence were included. Non-destaining boutons were defined as the boutons losing >10% of raw FM1-43 fluorescence only to 10Hz stimulation. All measurements were conducted on raw, unaltered images acquired using the same hardware configuration throughout the study. The total loss of fluorescence of each terminal during activity-dependent destaining was computed from individual puncta by standardizing the raw fluorescence to a common value for every bouton in the dataset, and then subtracting the averaged fluorescence after the 10Hz stimulus train from the averaged fluorescence of before 2Hz electrical stimulation. Since the FM1-43 loading procedure was consistent for slices from all treatment groups, the total amount of FM1-43 fluorescence in each bouton was assumed to approximate the number of releasable vesicles at the specific release site, as previously described ^88^. The release kinetics for each bouton during 2Hz electrical stimulation were determined from the time constant (*τ*) of an exponential decay function fitted to the FM1-43 fluorescence intensity during FM1-43 destaining. If the fit resulted in an R2 < 0.6, the ROI was excluded, providing an objective selection criterion for boutons displaying clear stimulus-dependent FM1-43 destaining. The rate of FM1-43 destaining, t_1/2_, was then computed by multiplying *τ* with 0.69 ^89^. The median t_1/2_ from the boutons within an individual slice was treated as an independent observation, and group comparisons were conducted between slices from different treatment groups.

#### GCaMP imaging and analysis

Brain slices were prepared from D1Cre mice, at least three weeks after striatal stereotactic injections of AAV5 virus transducing *cre-*dependent GCaMP6f or GCaMP7s expression in the striatonigral neurons and their axon terminals the SNr. After recovery at room temperature, the slices were transferred to the recording chamber on the stage of the two-photon microscope and was continuously superfused with aCSF. Imaging of presynaptic Ca^2+^ transients in striatonigral terminals was performed during bipolar electrical stimulation with stimulus trains of 2-10 Hz, lasting 50 s. After a first set of stimuli, DA (with 50 µM metabisulfite) or D1 receptor agonists were dissolved in the aCSF, and the slices were perfused with the solution for 5-10 min. Subsequently, electrical stimulation was applied in drug-containing aCSF. Thus, each fluorescent spot with an evoked Ca^2+^ transient could be examined in relation to drug-free conditions. In some experiments, the superfusion was then switched to drug-free aCSF and the effect of electrical stimulation was re-evaluated to determine if the drug’s effects could be reversed.

Images (16-bit, 512 x 512, pixel size 0.069 µm/pixel) were acquired from 80-100 µm into the slice, with a sampling frequency of 1 Hz using ScanImage version 2021.0.0.3baa4b74be. The time-series were imported into Fiji, where they were registered with Stackreg, and multiple circular ROIs per image were manually delineated around regions exhibiting GCaMP activity during 10Hz stimulation. While fluorescent elements, mostly fibers, were present in the absence of electrical stimulation, they remained unchanged upon exposure to electrical stimulation, chemical depolarization with 20 mM K^+^, or DA. These structures are likely fibers that have been severed during brain slice preparation, resulting in the loss of the Ca^2+^ buffering capacity that maintains a low GCaMP signal in intact axons. The mean fluorescence intensity was measured for each ROI across the time-series. For each ROI, the average intensity of the first 10 image frames was calculated, and then each frame’s intensity value was divided by this average to compute normalized values relative to the background control before stimulation (F_0_). The normalized values of the ROIs were then averaged per slice. The relative increase (F/F_0_) was used to compare between treatment groups, calculated by averaging the intensity of 10 image frames during electrical stimulation.

### Immunostaining and confocal microscopy

Mice were deeply anesthetized with pentobarbital (100 mg/kg, i.p) and transcardially perfused with ice-cold 4% (w/v) paraformaldehyde in 0.1 M sodium phosphate buffer (PFA, pH 7.5). The brains were dissected and immersed in the same fixative over night at 4°C. Using a vibrating microtome (Leica), 30 µm thick sections were cut through the striatum and the midbrain and were stored at -20°C in cryoprotectant solution containing (vol/vol) 30% ethylene glycol, 30% glycerol and 0.1 M sodium phosphate buffer (pH 7.5), until processed for immunofluorescence. Sections were washed three times for 10 min in Tris-buffered saline (50 mM Tris, 150 mM NaCl, pH 7.5; TBS) and then incubated for 1h in TBS containing 10% normal goat serum (NGS) and 0.3% Triton X-100. The sections were then incubated overnight at 4°C with gentle agitation in primary antibodies diluted in TBS containing 2% NGS and 0.3% Triton-X100. The primary antibodies were a rabbit anti-DARPP-32 (Cell signaling Technologies, 2306) diluted 1:500 and a chicken anti-GFP (Abcam, Ab13970) diluted 1:2000. The following day, after another three 10 min rinses in TBS, the sections were put in TBS containing goat-raised fluorescently conjugated Alexa 488, Alexa 568, Alex 647 (Life Technologies) secondary antibodies, and 2% NGS and 0.3% Triton-X100, for 45 min at room-temperature. After three washes in TBS the sections were mounted onto Superfrost microscope slides (Fisher Scientific) and Fluoromount-G anti-fading medium containing DAPI (SouthernBiotech).

Images were capture with a Zeiss LSM800 confocal system equipped with argon, DPSS and He/Ne lasers, and the Zeiss Blue software. In double-labeling experiments, the fluorophores were imaged sequentially with the detectors adjusted to prevent bleed-through between different channels. Images were acquired with a 20x/0.7NA air objective in 2048 × 2048 or 512 × 512 resolution (pixel size = 94.6 - 378 nm) and single frames were averaged 4 times. The number of immunopositive cells with staining above background (staining in corpus callosum) were counted using analyze particles feature of Fiji.

### Drugs

All drugs were dissolved as stock solutions at saturating concentrations on the day of experiments, in the vehicle suggested by the manufacturer. Aliquots of stock solution was diluted directly into the superfusing aCSF. DA (3,4-Dihydroxyphenethylamine hydrochloride, Sigma) was diluted to the final concentration aCSF right before application to the slice. The aCSF for DA experiments contained 50 µM sodium metabisulfite (Sigma) to delay oxidation. The following substances were purchased from Tocris, dissolved in DMSO and applied to the slice at the indicated concentrations: 1 µM SCH39166, ((6aS-trans)-11-Chloro-6,6a,7,8,9,13b-hexahydro-7-methyl-5H-benzo[d]naphth[2,1-b]azepin-12-olhydrobromide), 1 µM sulpiride ((RS)-(±)-5-Aminosulfonyl-N-[(1-ethyl-2-pyrrolidinyl)methyl]-2-methoxybenzamide), 100 µM picrotoxin, 5 µM NAS-181 ((2R)-2-[[[3-(4-Morpholinylmethyl)-2H-1-benzopyran-8-yl]oxy]methyl]morpholine dimethanesulfonate). The maximal concentration of DMSO in aCSF was 0.1% (w/w), which did not produce a discernable difference from DMSO-free aCSF in any of the experiments. The following drugs, also from Tocris, were dissolved in water before dilution in aCSF at the indicated concentrations: 10 µM amphetamine ((+)-α-Methylphenethylamine hemisulfate salt), 1µM SKF83566 (8-Bromo-2,3,4,5-tetrahydro-3-methyl-5-phenyl-1H-3-benzazepin-7-ol hydrobromide), 1-10 µM SKF81297 ((±)-6-Chloro-2,3,4,5-tetrahydro-1-phenyl-1H-3-benzazepine hydrobromide), 1-10 µM SKF38393 ((±)-1-Phenyl-2,3,4,5-tetrahydro-(1H)-3-benzazepine-7,8-diol hydrobromide), 10µM NKH477 (N,N-Dimethyl-(3R,4aR,5S,6aS,10S,10aR,10bS)-5-(acetyloxy)-3-ethenyldodecahydro-10,10b-dihydroxy-3,4a,7,7,10a-pentamethyl-1-oxo-1H-naphtho[2,1-b]pyran-6-yl ester β-alanine hydrochloride), 1 µM quinpirole ((4aR-trans)-4,4a,5,6,7,8,8a,9-Octahydro-5-propyl-1H-pyrazolo[3,4-g]quinoline hydrochloride), 1 µM SCH23390 ((R)-(+)-7-Chloro-8-hydroxy-3-methyl-1-phenyl-2,3,4,5-tetrahydro-1H-3-benzazepine hydrochloride). TTX (tetrodotoxin citrate) was purchased from HelloBio, dissolved in water, and applied at 1 µM concentration in the aCSF.

## Acknowledgments

We extend our gratitude to Gilad Silberberg, Abdel El Manira, Gilberto Fisone, Konstantinos Meletis, Alessandro Usiello, Sten Grillner, Peter Löw, Ole Kiehn and all the members of the Fisone, Borgkvist and Santini labs for their invaluable methodological assistance and insightful discussions, and to Kristoffer Tenebro Berglund, the staff, and the veterinarians of the Comparative Medicine Biomedicum (KM-B) for their continuous support and assistance with the maintenance of the mouse colonies. This work was supported by the Knut and Alice Wallenberg Foundation (Wallenberg Academy Fellow Grant KAW 2017-0169 and project grant 2020-0054 to E.S.), the Swedish Research Council (2016-02758 to E.S. and 2016-03129 to A.B.), the Stiftelsen Olle Engkvist Byggmästare (E.S. and A.B.), Åhlén’s foundation (A.B.), Magn. Bergvall’s foundation (A.B.), The Strategic Research Program in Neuroscience (StratNeuro) starting (E.S. and A.B.) and bridging (E.S.) grants, Karolinska Institute starting (E.S.) and KID (E.S. and J.C.R.) grants. D.S. and is supported by NIDA R01DA07418 and JPB Foundation. O.J.L. was supported by NIH F30 MH114390.

## Author contributions

Conceptualization, A.B., E.S. and D.S; Methodology, M.M., A.B. and E.S.; Validation, A.B. and E.S.; Formal Analysis, A.B. and E.S.; Investigation, M.M., O.J.L. and A.B.; Resources, A.B., E.S. and D.S.; Writing - Original Draft, A.B. and E.S.; Writing - Review & Editing, M.M, O.J.L., D.S., A.B. and E.S.; Visualization, A.B.; Supervision, A.B. and E.S.; Project Administration, A.B. and E.S.; Funding Acquisition, A.B., E.S. and D.S.

